# Single-oocyte transcriptional profile of early-stage human oocytes reveals differentially expressed genes in the primordial and transitioning stages

**DOI:** 10.1101/2025.07.01.662667

**Authors:** J. H. Machlin, D. F. Hannum, A. S.K. Jones, T. Schissel, K. Potocsky, E. E. Marsh, S. Hammoud, V. Padmanabhan, J.Z. Li, A. Shikanov

## Abstract

The critical initial step in human oocyte maturation - the transition of ovarian follicles from dormancy to activation - remains poorly understood. Here we performed RNA sequencing on single oocytes isolated from early-stage follicles from nine healthy reproductive-age donors. Data for 133 high-quality oocytes formed two connected clusters, C1 and C2, with 5,449 significantly differentially expressed genes. Using recently reported gene lists for early-stage follicles we found that C1 oocytes likely came from earlier, dormant primordial follicles, while C2 oocytes match later-stage primordial or transitioning follicles. We sought to validate two DE genes, UHRF1 for C1 and CCN2 for C2, by their RNA-FISH in situ pattern in morphologically classified follicles, but did not observe statistically significant differences between follicle stages. This apparent discrepancy between follicle’s stage determined by its morphology and oocyte’s transcriptional state, if replicated in additional studies, may indicate a lack of closely coupled synchrony between follicle morphology and oocyte’s functional state.

## Introduction

Ovarian follicles are the source of both fertility and endocrine function. Follicles contain an oocyte surrounded by hormone producing somatic cells that grow in a coordinated process known as folliculogenesis. During this process, dormant oocytes activate, begin transcribing and storing mRNAs in preparation for fertilization, and undergo tremendous volumetric expansion, growing from ∼20 µm in diameter to ∼120 µm in humans^1–4^. The critical initial step in oocyte maturation, transitioning from dormancy to activation, remains poorly understood. This lack of knowledge has hindered progress in activating and maturing human primordial follicles in culture, which is of interest for clinical interventions. A significant number of genetic knockout experiments have been conducted to identify key signaling pathways and transcription factors involved in follicle activation and transcriptional changes in transitioning oocytes ^5–7^. However, these pathways may not be conserved in lower species when compared to humans, leaving a large gap in our understanding^8^.

Recently, advances in single-cell RNA-sequencing technology provided an opportunity to explore the cellular heterogeneity and unique transcriptional shifts that arise during folliculogenesis^9^. Two recent studies have performed the laborious task of sequencing individual mouse oocytes by first isolating follicles and then removing the somatic cells. These studies identified transcriptional signatures of oocytes transitioning from the primordial to primary stage in mouse ovaries, providing a valuable set of genes for reference^10,11^. However, very few studies have investigated the transcriptional profiles and the primordial-primary transition in early-stage human oocytes, due to the rarity of tissue from healthy reproductive-age individuals and the technical complexity involved in efficiently removing early-stage follicles from the ovarian cortex. For example, analyses of human ovaries by Wagner et al. and Zhang et al. relied on donors who had previously undergone androgen therapy or from cancer patients, focusing their analysis on oogonial stem cells and interactions between granulosa cells and oocytes^12,13^. In a more recent study, Rooda et al. investigated transcriptional differences during follicle activation by using isolated follicles at various stages and sequencing each follicle without separating the oocytes from the surrounding somatic cells^14^, staging the follicles by using existing classification criteria based on granulosa cell morphology (squamous, cuboidal, or both)^15^. While the somatic contribution could not be decoupled from the oocyte transcriptome, Rooda et al. identified unique transcriptional signatures for follicles at different activation stages, thus providing a new set of reference gene lists for follicle stages. Here, we focused on transcriptional changes in human oocytes isolated from early-stage follicles and separated from somatic cells.

To accomplish this, we isolated and sequenced 216 oocytes from the ovarian cortex of nine healthy reproductive-age donors. We developed a protocol to specifically enrich for follicles <70µm and efficiently remove their somatic cells to obtain the denuded oocytes while maintaining oocyte viability. The single-oocyte transcriptomes formed two interconnected clusters with distinct gene signatures. By using the recently published gene sets distinguishing follicle stages^14^ we showed that the two clusters correspond to “earlier-stage” and “later-stage” oocytes, respectively. Our analysis revealed key biological processes associated with these oocyte stages and may further inform the development of *in vitro* activation (IVA) protocols.

## Methods

### Collection of Human Ovarian Tissue

This study used tissue samples from ten de-identified deceased donors, nine were used for single-cell RNA-sequencing analysis and one additional donor was used for validation using RNA FlSH. This tissue was procured through the International Institute for the Advancement of Medicine (IIAM) and the associated Organ Procurement Organization (OPO) involved in the harvest. All ten donors were pre-menopausal and examination of provided medical records indicated no pathological conditions impacting ovarian function (for their age, BMI, recorded “race” and Cold Ischemic Time, see **Table I** and **Supplemental Table 1**). Cold ischemic time (CIT) was calculated as the time interval between cross-clamp time of the donor (and subsequent cessation of arterial blood flow to the ovaries) in the operation room and start time of the tissue harvest after arrival at the laboratory. Before cross-clamp, the organs were perfused with Belzer University of Wisconsin® Cold Storage Solution (Bridge of Life, SC, USA), Custodiol® HTK (Histidine-Tryptophan-Ketoglutarate) Solution (Essential Pharmaceuticals, NC, USA), or SPS-1 Static Preservation Solution (Organ Recover Systems, IL, USA).

### Ethical Approval Process

The IIAM procures tissue and organs for non-clinical research from Organ Procurement Organizations (OPOs), which comply with state Uniform Anatomical Gift Acts (UAGA) and are certified and regulated by the Centers for Medicare and Medicaid Services (CMS). These OPOs are members of the Organ Procurement and Transplantation Network (OPTN) and the United Network for Organ Sharing (UNOS) and operate under a set of standards established by the Association of Organ Procurement Organizations (AOPO) and UNOS. Informed, written consent from the deceased donor’s family was obtained for the tissue used in this publication. A biomaterial transfer agreement is in place between IIAM and the University of Michigan that restricts the use of the tissue for pre-clinical research that does not involve the fertilization of gametes. The use of deceased donor ovarian tissue in this research is categorized as ‘not regulated’, per 45 CFR 46.102 and the ‘Common Rule’, as it does not involve human subjects and complies with the University of Michigan’s IRB requirements as such.

### Tissue Processing

All tissue processing was done aseptically in a biosafety cabinet. After receiving donor tissues, the ovaries were separated from the uterus and fallopian tubes. The ovaries were decortified using a custom cutting guide (Reprolife Japan, Tokyo) to remove approximately 1mm thick cortex pieces that were approximately 10mmx10mm squares. The squares were then aseptically transferred into holding media (Quinn’s Advantage Medium with HEPES (QAMH), 10% Quinn’s Advantage Serum Protein Substitute (SPS), CooperSurgical, Måløv, Denmark) before cryopreservation.

### Slow Freezing Procedure

The methods as described by Xu et al. were used for slow freezing^16^. Briefly, squares of cortical tissue approximately 10mm x 10mm x 1mm were placed into cryovials (Nunc, Roskilde, Denmark) filled with pre-cooled cryoprotectant media (QAMH, 10% SPS, 0.75M dimethyl sulfoxide (DMSO) (Sigma Aldrich, St. Louis, USA), 0.75M ethylene glycol (Sigma Aldrich, St. Louis, USA), 0.1M sucrose (Sigma Aldrich, St. Louis, USA)), and equilibrated at 4°C for at least 30 minutes. After equilibration, cryovials were loaded into the Cryologic Freeze Control System (Cryologic, Victoria, Australia). Vials were then frozen using the following protocol: (1) cooled from 4°C to-9°C at a rate of-2°C /min (2) equilibrated for 6 min at-9°C (3) seeded manually using large swabs cooled by submersion in liquid nitrogen (4) held for 4 min at-9°C (5) cooled to-40°C at a rate of - 0.3°C/min and (6) plunged into liquid nitrogen and stored in a cryogenic storage dewar until thawed for use.

### Tissue Thawing Procedure

Vials with ovarian tissues were removed from liquid nitrogen and placed in a 37°C bath. Once the cryoprotectant media in the vial thawed, the tissue was transferred into “Thaw Solution One” (1M DMSO, 0.1M Sucrose, 10% SPS in QAMH) for ten minutes. Tissue was then incubated sequentially in “Thaw Solution Two” (0.5M DMSO, 0.1M Sucrose, 10% SPS in QAMH), “Three” (0.1M Sucrose, 10% SPS in QAMH), and “Four” (10% SPS in QAMH) for ten minutes each. Thaw solutions were kept at room temperature, and samples were protected from light and agitated while in thaw solutions.

### Tissue Dissociation and Collection of Oocytes for scRNAseq

Slow frozen/thawed ovarian cortex squares 10mm x 10mm x 1mm were dissected into ∼1 mm cubes at room temperature in Thaw Solution Four using a McIlwain Tissue Chopper (Ted Pella, Inc., Redding, CA). The tissue cubes were rinsed twice with Dulbecco’s Phosphate Buffered Saline without calcium or magnesium (DPBS^-/-^) (Fisher Scientific, US), then transferred to a digestion solution containing 1 mg/mL Collagenase IA (Sigma Aldrich, Germany) and 0.02 mg/mL DNase I (Worthington Biochemical, US) in DPBS with calcium and magnesium (DPBS^+/+^, Fisher Scientific). Tissue was transferred to a shaker at 150 rpm to digest for 2 hours at 37°C, then the tissue was strained through a 70 μm strainer (Fisher Scientific), and the cell suspension was quenched with ice cold 10% fetal bovine serum (FBS, Fisher Scientific) in DBPS^-/-^. The quenched cell suspension was strained again through a 30 μm strainer (PluriSelect, US), the strainer was then backwashed into a clean 30 mm dish using DBPS^-/-^. This backwashed cell suspension was divided between several 30 mm dishes and follicles between 70 μm and 30 μm were manually collected using The Stripper Micropipetter (CooperSurgical, Denmark) with a 75 µm stripper tip and placed in 45 μL droplets of Leibovitz’s L-15 medium (L15) (Fisher Scientific) under Vitrolife Inc Ovoil (Fisher Scientific). All collected follicles were transferred for a second digest in solution containing 1 mg/mL Collagenase IA (Sigma Aldrich) and 0.02 mg/mL DNase I (Worthington Biochemical) in DPBS^+/+^ and transferred to a shaker at 150 rpm to digest for 1 hour at 37°C. The digest was quenched with ice cold 10% fetal bovine serum (FBS, Fisher Scientific) in DBPS^-/-^ and separated into 4-well plates for follicle collection. Follicles and denuded oocytes were manually collected using The Stripper Micropipetter (CooperSurgical) with a 75 μm stripper tip and placed in 45 μL droplets of Leibovitz’s L-15 medium (L15) (Fisher Scientific) under Vitrolife Inc Ovoil (Fisher Scientific). Follicles were gently pipetted up and down to remove surrounding cells and release oocytes. Denuded oocytes were placed individually into wells of a V-bottom 96-well plate containing 7 μL of lysis buffer (seqWell Inc. Beverly, MA) and the plate was placed at-80°C until ready for sequencing.

### Histological Analysis

Fresh cortex strips were fixed in Bouin’s fixative overnight at 4°C, washed three times with DPBS^-/-^ and stored in 70% ethanol. All samples were processed at the Histology Core in the Dental School at the University of Michigan. Fixed tissue was embedded in paraffin blocks, serially sectioned at a thickness of 5µm with 4 sections per slide up to 50 slides. Every other slide was stained with hematoxylin and eosin (H&E). The H&E stained slides were sent to the Unit for Laboratory Animal Medicine (ULAM) Core at the University of Michigan where they were whole slide scanned at 20x on an Aperio digital whole slide scanner (Aperio AT2). Follicles were counted and staged in one section on each slide, skipping approximately 40µm between the sections (**Figure 1** and **Supplemental Figure 1**). All primordial, transitional primordial and primary follicles were counted in each analyzed section. All larger follicles were only counted where the nucleus was visible to avoid “double-counting”. Due to the ovarian tissue variability we counted 2-25 slides for the nine donors, equating to an average tissue volume of 3.1 mm^3^ with a standard deviation of 1.7 mm^3^. In order to obtain a density measurement, the area was calculated for all tissue sections counted using FIJI (ImageJ). Follicle densities for each section counted were calculated using the area measurement multiplied by the thickness of the tissue between the sections we counted and plotted using GraphPad Prism software. Follicles were staged based on standard morphological nomenclature previously described^17^.

**Figure 1.**
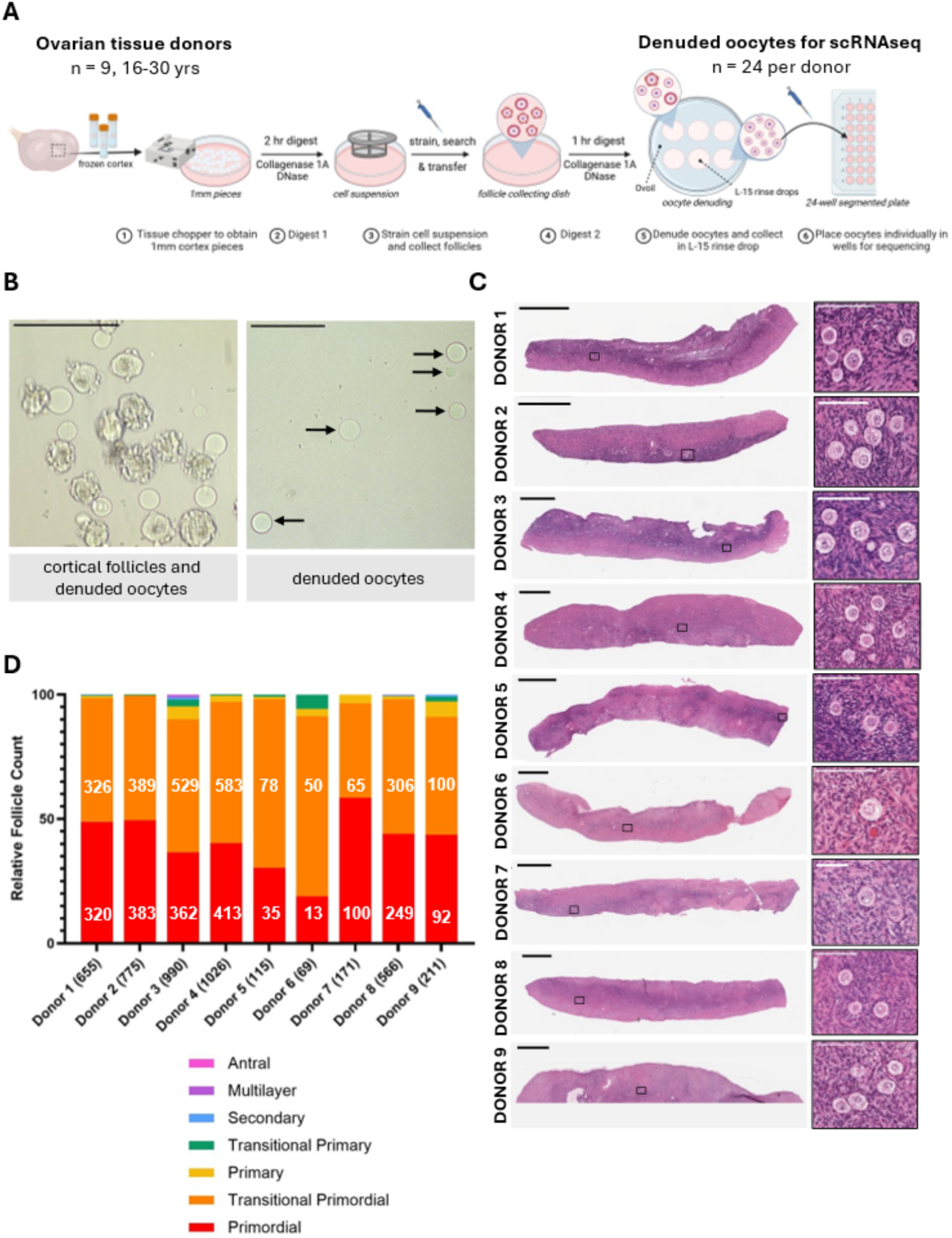
Experimental process and tissue histology. **A.** Schematic summary of the ovarian cortical tissue dissociation and single-oocyte collection process. Human ovarian tissue from 9 donors ranging from 16-30 yrs was used and, for each donor, 24 denuded oocytes from follicles within the ovarian cortex were collected for scRNAseq. **B.** Bright-field images of examples of dissociated cortical follicles (left) and the subsequently denuded oocytes (right), as indicated by arrows. Scale Bars (SB): 100µm. **C.** H&E images of ovarian cortex strips (left) and zoomed-in view of follicles (right) for the 9 donors in this study. Strip SB: 1mm. Zoomed-in SB: 100µm. **D.** Relative proportion of follicles across follicle stages in the ovarian cortex, compared across the 9 donors. The total number of follicles counted for each donor is indicated in the x axis label. The number of follicles for the primordial and transitioning primordial stages are shown in their corresponding segments in the bar plot. Created with BioRender.com.

### Sequencing Data Collection

We isolated 24 oocytes for RNAseq analysis from each of the nine donors. Oocytes were moved into lysis buffer in 96-well plates, stored at-20°C, and submitted on dry ice to the Advanced Genomics Core at the University of Michigan. RNAseq library preparation used the PlexWell scRNA reagents (seqWell Inc. Beverly, MA) according to the manufacturer’s instructions. The Core assessed the final library quality using the LabChip GX (PerkinElmer, Hopkinton, MA), which measures nucleic acid fragment size. Pooled libraries were subjected to 150 bp paired-end sequencing on the NovaSeq6000 (Illumina, San Diego, CA). Sequence reads in FASTQ files were aligned to GRCh38 (ENSEMBL), using STAR v2.7.8a, resulting in a cell-by-gene count matrix for 60,628 genes and the 216 samples. From the raw counts data we calculated the mean count for each gene, and found 18,178 genes with mean >1. Subsequent analysis used these 18,178 genes.

### Data Analysis

Data processing and analysis was performed with R (v4.3.3), and most plotting was done with ggplot2 (3.5.0). From the cell-by-gene count matrix, samples were filtered to retain those with (1) number of transcripts (i.e., “library size”) > 500,000 and (2) percentage of reads coming from mitochondria-coded RNAs < 20%, resulting in 165 samples out of the 216. We then normalized the count data for each sample to transcripts-per-million, adding a floor value of 0.2 (to make the 0 counts amenable for taking the log), then converted to log(base=2) values. Further processing was done using Seurat (v5.0.3), including finding the 2,000 most highly variable genes (HVGs) using the “vst” method, scaling data by gene, and running PCA using the 2000 HVGs. As PC1-2 patterns were largely driven by library size (not shown), we further selected samples with PC1> (-40) and PC2<12, resulting in 133 samples (**Table 1**). We used the top 10 PCs to obtain the UMAP projections (**Figure 2A**) and identify 2 clusters at resolution=0.4, with 48 and 85 samples in Cluster 1 and 2, respectively. Differential gene expression analysis between Cluster 1 and 2 samples was done in Seurat using the default Wilcoxon-Rank Sum test. Gene-level results were included in **Supplemental Data File S1 and S2** and shown in **Figure 2**. We submitted the fold-change and FDR results to *LRPath* (online version on Nov 18, 2024) to perform pathway enrichment analysis across GO biological processes, GO cellular component, GO molecular function, KEGG Pathway, Panther Pathways, and MeSH terms. Of the 18,178 genes, only 14,895 had Entrez IDs, of which another 138 were not recognized by LRPath. We focused the analysis of enrichment results on GOBP terms (**Supplemental Data File S3** and **Figure 2C**).

**Figure 2.**
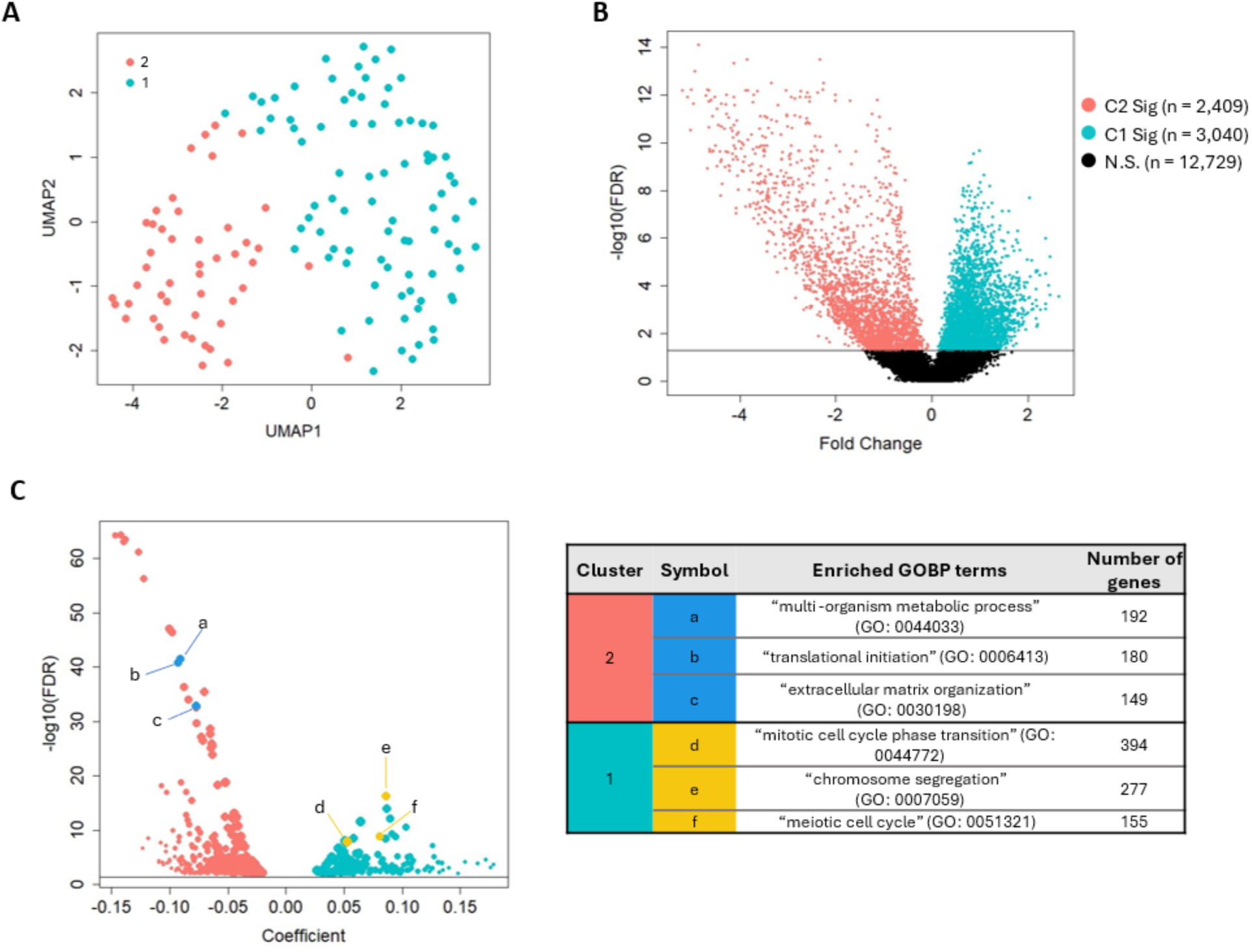
Oocytes from human ovarian cortex follicles show two interconnected clusters. **A.** Uniform manifold approximation and projection (UMAP) visualization of RNA expression data for 133 pass-QC oocytes showed two interconnected clusters, with 48 and 85 oocytes in Cluster 2 and Cluster 1, respectively. **B.** Differential expression (DE) analysis revealed 2,409 and 3,040 genes to be significantly higher in Cluster 2 and Cluster 1 oocytes, respectively. The Volcano plot of genes is shown. **C.** Gene ontology (GO) enrichment analysis of DE genes for GO-Biological Processes (GOBP) revealed significantly enriched terms in their Volcano plot (left). Examples of highly enriched terms, A-C for genes expressed higher in Cluster 2 and D-F for those higher in Cluster 1, are shown in the table (right).

**Table 1.**
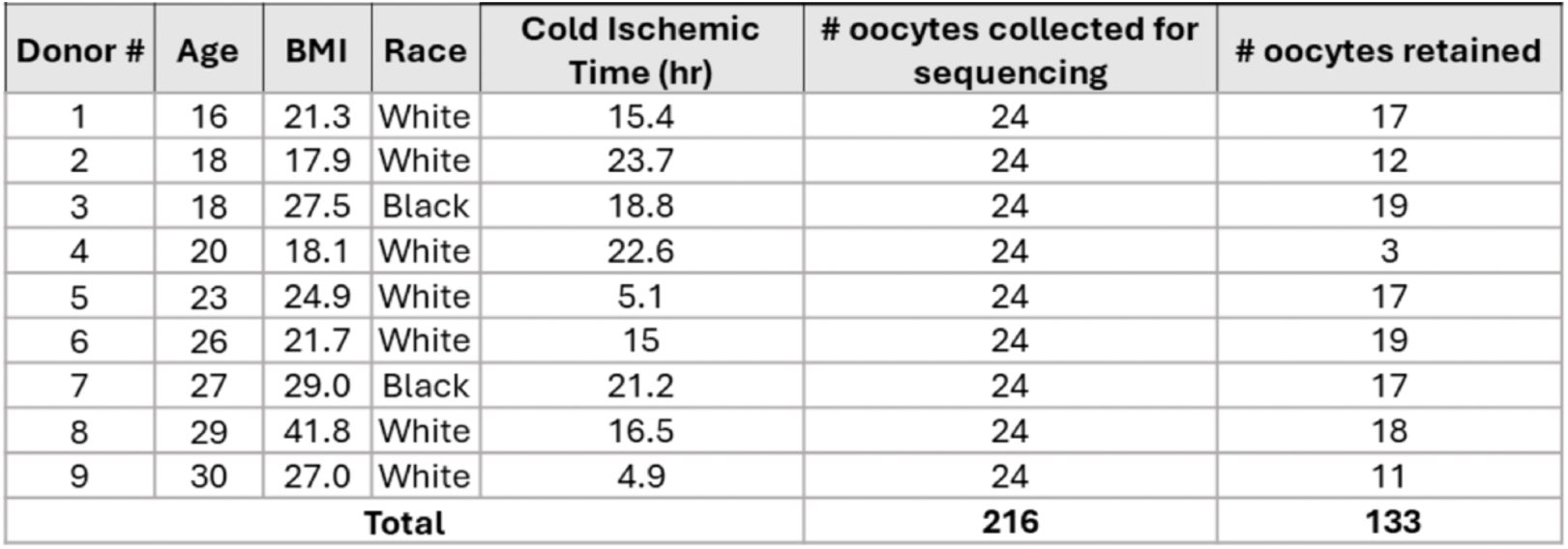
Donor metadata, including age. BMI, reported race, Cold ischemic Time of the ovary sample, number of oocytes collected for sequencing analysis, and the number passing quality control and used in data analyses.

### Comparative analyses with DEGs from Rooda et al

We downloaded several sets of DE genes from the supplementary data of Rooda et al.^14^: SU4A, for genes that are high in the primordial, transitioning, and primary stages; and SU11, for comparison between the dormant and active subgroups of their primordial follicles. Specifically, SU4A contained T-scores for differentially expressed genes – in both directions – for three of the relevant comparisons, t1: primordial vs. intermediate, t2: primordial vs. primary, and (t3) intermediate vs. primary. We defined “primordial-high” genes as those with t1>0 and t2>0; “intermediate-high” genes as those with t1<0 and t3>0; and “primary-high” genes as those with t2<0 and t3<0. Similarly, we used T-scores in SU11 to define “active-high” and “dormant-higher” genes.

The number of these three sets of genes, and those overlapping with our DE genes, was reported in **Figure 3A** and **Figure 4A**, and the lists of genes can be found in **Supplemental Data File S4** and **S5** respectively.

**Figure 3.**
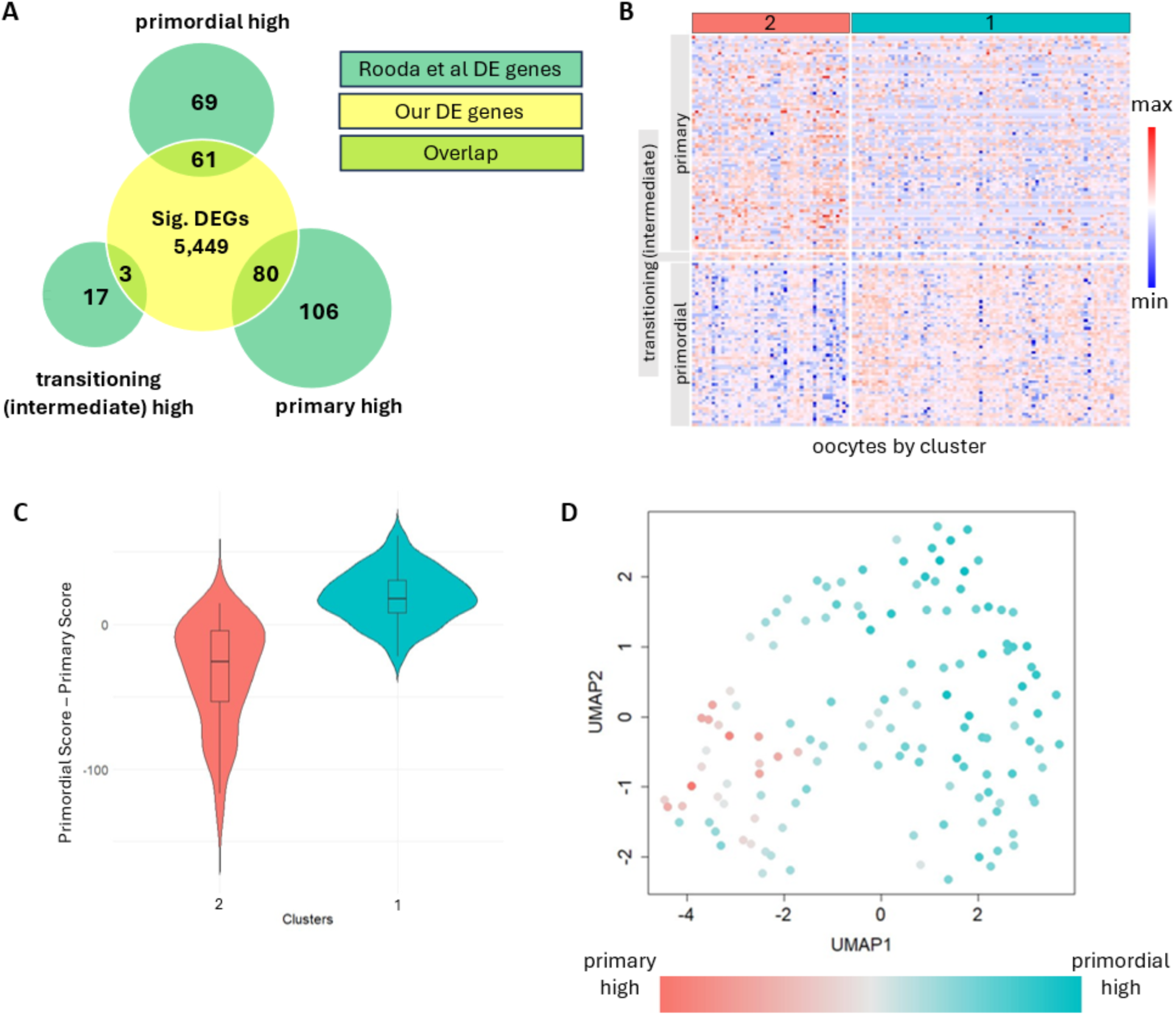
Dataset integration reveals oocyte cluster identities as corresponding to early-later stage transitions. **A.** Comparison of our single-oocyte DEGs to Rooda et al. DEGs expressed higher in primordial, transitioning (intermediate) and primary follicles finds 61, 3, and 80 overlapping genes for the three stages, respectively. **B.** Expression pattern of the 61, 3, and 80 overlapped DEGs across our 133 oocytes, ordered as Cluster 2-left and Cluster 1-right, shows a higher expression of primary genes in Cluster 2 and a higher expression of primordial genes in Cluster 1. **C.** Violin plot of per-oocyte primordial-score-minus-primary-score, calculated by summing each oocyte’s log expression values for the 61 and 83 genes in B, and compared between Cluster 1 and Cluster 2. **D.** Overlay of the oocytes’ primordial-score-minus-primary-score as symbol color in the 133-oocyte UMAP projection, previously shown in Fig. 2A, confirms the primordial-to-primary gradient in the transition from Cluster 1 to Cluster 2.

**Figure 4.**
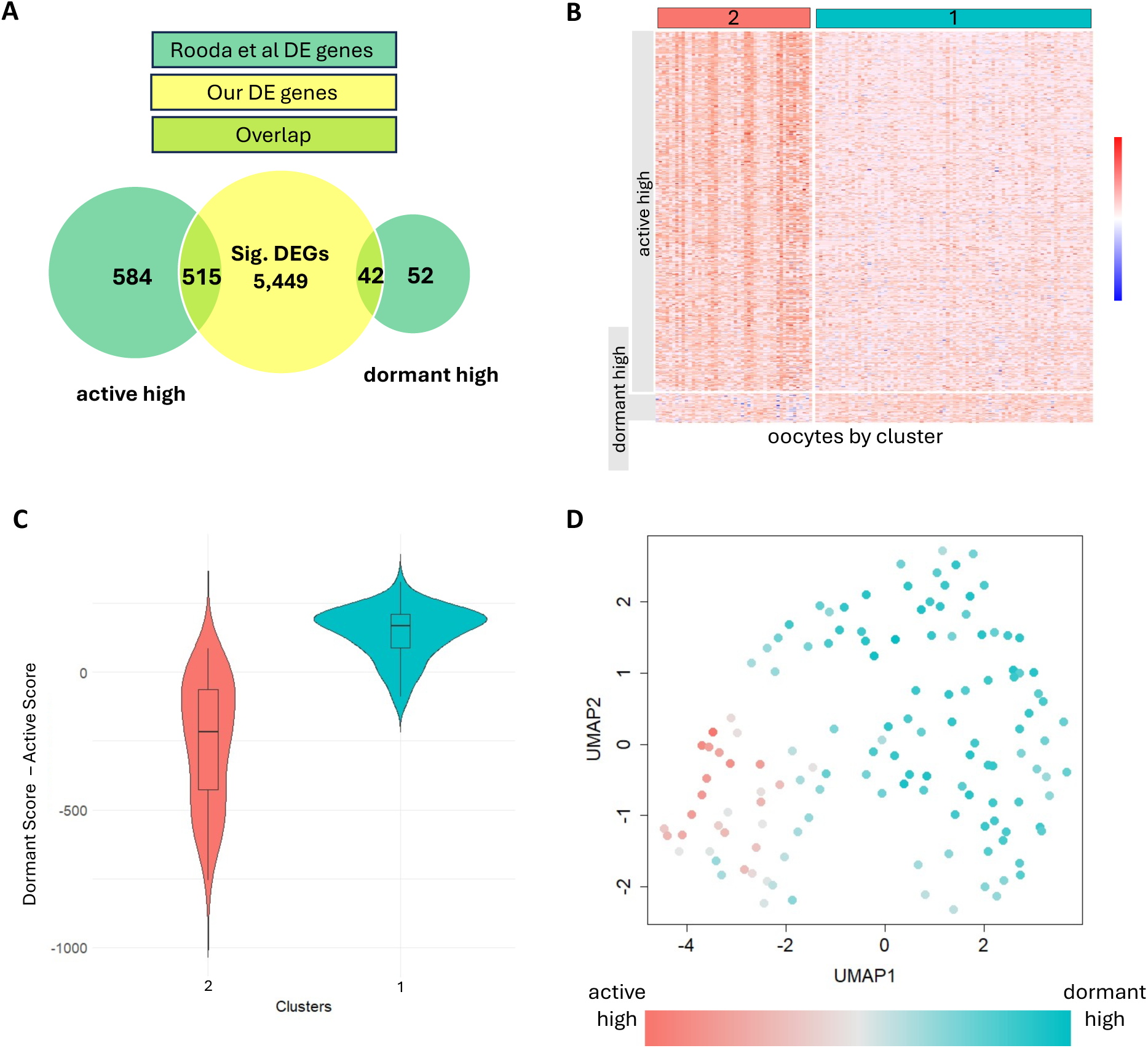
Oocyte cluster identities align with the reported dormant-to-active transition in follicles. **A.** Comparison of our single-oocyte DEGs to Rooda et al. DEGs in their active-dormant follicle comparisons shows overlapped genes with our dataset. **B.** Expression patterns of the 515 active-high and 42 dormant-high overlapped genes across 133 oocytes in our data, ordered by Cluster 2-left and Cluster 1-right. **C.** Violin plot of the oocytes’ dormant-score-minus-active-score, calculated by summing log expression values for the 42 and 515 genes, respectively, and compared between Cluster 1 and Cluster 2. **D.** Alignment of the dormant-to-active gradient with the transition from Cluster 1 to Cluster 2, shown in the original UMAP projection with symbol color indicating the oocytes’ dormant-score-minus-active-score.

From our scaled log expression data, we extracted the values for the overlapped genes for all 133 oocytes, and displayed the heatmap in **Figure 3B** and **Figure 4B**. Here scaling is for each gene, so that the gene’s expression values across the 133 oocytes have mean=0 and sd=1. To calculate each oocyte’s overall activity score for the primordial, primary (combined with the 3 genes for transitioning), dormant, and active gene sets, we summed the scaled log expression values for the genes in that gene set, and then constructed the “delta”, which is the primordial-primary scores, or dormant-active scores, and compared between the two clusters in **Figure 3C** and **Figure 4C**, and showed the color-coded UMAP **in Figure 3D** and **Figure 4D**.

### RNA Fluorescence In Situ Hybridization (FISH)

Slides were stored at room temperature in a slide box containing a silica desiccator packet prior to staining. Slides were baked for 1 hour in a 60°C dry oven the night before staining. RNA FISH probes were acquired from Advanced Cell Diagnostics (ACD) and stained using the RNAscope multiplex fluorescent manual protocol and kit (RNAscope Multiplex Fluorescent Reagent Kit v2, ACD, Newark, CA). The RNAscope Probes Hs-CTGF (aka *CCN2*) (ACD catalog 560581) and Hs-UHRF1 (ACD catalog 559901) were used. The RNA FISH protocol was performed according to the manufacturer’s instructions (ACD document number 323100-USM) with a 15-minute antigen retrieval and a 45-minute protease treatment. Slides were counterstained with Hoechst 33342 (Fisher Scientific) for 30 minutes at room temperature, and mounted on microscope slides with Prolong Diamond (Fisher Scientific). Experiments for *CCN2* and *UHRF1* quantification were performed using ovarian tissue from Donor 10.

### Confocal microscopy and quantification of CCN2 and UHRF1 expression

Images were captured on a Leica SP8 laser scanning confocal microscope. All confocal images were captured using identical laser power, gain, and imaging parameters for each probe. Follicles were imaged at the cross-section where the oocyte nucleus was clearly visible, and a z-stack was set up from the top visible plane of the follicle to the bottom visible plane of the follicle. All RNA FISH quantification was performed in FIJI^18^ where on unaltered images that contained metadata. To quantify the expression of *CCN2* and *UHRF1* in oocytes, an ROI was created around the oocytes at the center z stack slice for each image. Images were then subjected to threshold and watershed. Individual mRNA puncta were counted using the “analyze particles” feature in FIJI. The individual mRNA puncta were reported for oocytes at the primordial, transitioning primordial, and primary stages.

### Statistical analysis

Statistical analysis was performed using GraphPad Prism 10. Graphs showing mRNA puncta counts were plotted as mean ± SD and statistical significance was determined using an unpaired t test (*p* < 0.05).

## Results

### Histological analysis of ovarian cortex tissue confirms presence of primordial, transitioning primordial and primary follicles for transcriptional profiling

Ovarian tissue in this study originated from nine healthy donors ranging from 16 to 30 years of age and without known gynecological diseases (**Table 1** and **Supplemental Figure 1A,B**). Our goal was to analyze transcriptomic signatures in oocytes from primordial, transitioning primordial, and primary follicles, therefore, accurate staging of follicles was critical. To analyze the follicular makeup of each ovarian cortex sample we performed histological evaluations of morphology, follicle density, and stage distribution. The stroma and follicles appeared morphologically normal in all nine cortex samples (**Figure 1C**). Across the nine samples, we visually identified 4,578 follicles, and for each sample measured follicle count-per-volume and the follicle stage distribution (**Figure 1D, Supplemental Table 2,** and **Supplemental Figure 1C).** As expected, a majority of follicles were in the primordial and transitioning primordial stages, making up an average 41.2% and 54.4% of all follicles, respectively. All donors except Donor 2 also contained a small portion (0.1% - 6.2%) of primary follicles, for an average of 2.6% of the total. In comparison, transitioning primary, secondary, multilayer and antral follicles were significantly rarer, accounting for an average of 1.8%. In sum, morphological evaluations confirmed that the analyzed oocytes were isolated from a range of early-stage follicles, based on the currently accepted methods for classification^15^.

### Single-oocyte RNA-sequencing analysis reveals two interconnected clusters

We collected 24 denuded oocytes from each dissociated ovarian cortex for single-oocyte RNA-sequencing, for a total of 216 oocytes (**Figure 1A,B**). After quality control (see **Methods**), 133 oocytes were retained for downstream analysis (**Table 1**). Plotting the 133 transcriptomes using Uniform Manifold Approximation and Projection (UMAP) revealed two interconnected clusters, Cluster 1 (C1) and Cluster 2 (C2), with a gradual transition between them (**Figure 2A**). Between the two clusters there were 5,449 significantly differentially expressed genes (DEGs) with 3,040 significantly high in C1 and 2,409 significantly high in C2 at FDR < 0.5 (**Figure 2B** and **Supplemental Data File S1 and S2**). To identify cluster-specific biological processes we submitted the DE results to *LRPath*^19^ for gene ontology (GO) analysis (**Supplemental Data File S3**).

Genes higher in C1 were enriched for terms “mitotic cell cycle phase transition” (GO: 0044772, 394 genes), “chromosome segregation” (GO: 0007059, 277 genes), and “meiotic cell cycle” (GO: 0051321, 155 genes) (**Figure 2C**). Genes of interest within these terms included *H1FOO*, encoding an oocyte-specific histone expressed in murine and human primordial follicle oocytes through the 2-cell embryo stage, essential for the development of oocytes^20,21^. Another gene, *SLC26A8*, is an anion transporter previously reported to be exclusively expressed in the testes during spermatogenesis but may have a role in oocytes^22^, and *DMRTC2*, is expressed specifically in the germ cells of fetal mouse ovaries and plays an important role in normal germ cell development and meiotic initiation^23^.

In contrast, genes higher in C2 were enriched for a variety of processes, such as “multi-organism metabolic process” (GO: 0044033, 192 genes), “translational initiation” (GO: 0006413, 180 genes), and “extracellular matrix organization” (GO: 0030198, 149 genes) (**Figure 2C**). These terms contained an abundance of genes encoding ribosomal proteins (*RPS20, RPL18, RPL31, RPS5, RPL6, RPL3*, to name a few). It is well known that during their growth phase, oocytes accumulate ribosomes to support protein synthesis during meiotic maturation and after fertilization^24^. These enriched processes suggested that the isolated oocytes span a range of developmental stages, with C1 highlighted by meiosis and C2 by translation initiation and ECM remodeling.

### C1-C2 transition corresponds to the genes for “earlier” and “later” stages in the primordial-to-primary follicle transition

We further evaluated the DEGs of our two oocyte clusters against the gene lists for different stages of follicles recently published by Rooda et al.^14^. They performed scRNAseq on cells from individually-staged whole follicles isolated from the ovarian cortex, including 17 primordial, 33 transitioning primordial or “intermediate”^14^, and 42 primary follicles, and reported differentially expressed genes in pairwise comparisons between stages. We first examined their top DE genes for primordial, transitioning primordial, and primary follicles, focusing on those that overlapped with the significantly differentially expressed genes from our clusters (**Figure 3A**). Encouragingly, standardized expression levels of Rooda et al. genes across our 133 oocytes correlated well with the C1-C2 distinction: the 61 primordial genes showed higher expression in C1, whereas the 80 primary and 3 transitioning primordial genes from Rooda et. al showed higher expression in C2 (**Figure 3B**), suggesting that C1 is more primordial-like while C2 is more primary-like.

We then used these gene sets to create a predictive score for each oocyte to summarize their primary-versus-primordial characteristics. Since there were only three transitioning primordial follicle genes and they showed a similar expression pattern as the primary follicle genes, we combined them and used the 83 genes to calculate a “primary” score and likewise calculated a “primordial” score using the 61 primordial genes (**Methods**). For each oocyte, we subtracted its primary score from its primordial score and saw a higher delta score in C1 (**Figure 3C**), again suggesting that C1 oocytes are primordial-like. We also visualized the delta in the original UMAP projection, which shows an early to late-stage transcriptome shift from C1 to C2 (moving from right to left) (**Figure 3D**).

### C1-C2 difference also aligned with dormant-active gradient reported for primordial follicles

Next, we examined an additional DE gene list from Rooda et. al. for comparing activated with dormant primordial follicles. From the reported 584 active-high and 52 dormant-high genes, 515 active and 42 dormant genes overlapped with our DE genes (**Figure 4A**). Their standardized expression levels across our 133 oocytes showed that the 42 dormant-high genes are expressed higher in C1, whereas the 515 active-high genes were higher in C2, (**Figure 4B,C**), consistent with C1 being less active than C2. When we calculated the delta between the predicted “active” and “dormant” scores for each oocyte (**Methods**) and visualized on the UMAP projection, there was a clear shift of dormant-to-active characteristics from C1 to C2 oocytes (**Figure 4D**), in parallel to the trend seen in **Figure 3D**. The two sets of delta scores were concordant, as seen when we plotted the primordial-primary scores against the dormant-active score across the 133 oocytes (**Supplemental Figure 2**). The genes used for the two types of scores: 144 genes in Figure 3 and 557 genes in Figure 4, both originating from Rooda et. al., shared only seven genes in common. Yet the two gene sets describe two highly aligned gradients in our 133 oocytes, one spanning primordial-primary follicles, the other between the dormant and the active phases of primordial follicles.

### Correlation of morphological staging to the transcriptome-based markers using RNA FISH

Next, we performed RNA FISH staining of cortical ovarian tissue using two candidate markers selected from our DEGs, one for C1 (primordial-like, or dormant) and the other for C2 (primary-like or active) and assessed their spatial patterns among follicles staged using traditional granulosa cell morphology parameters^15^. The gene for Ubiquitin Like with PHD and Ring Finger Domains 1, *UHRF1*, was expressed significantly higher in C1, while the gene for Cellular Communication Network Factor 2, *CCN2*, was significantly higher in C2, as depicted in the volcano plot (**Supplemental Figure 3**). Other than DE evidence, they were also chosen for their known functions related to the GO terms identified in the pathway analysis (**Figure 2C**). *UHRF1* is involved in DNA repair and cell cycle regulation^25,26^, while *CCN2* is a signaling molecule involved in wound healing and interactions between an activated follicle with its extracellular matrix components. Of note, *CCN2* is a gene target of YAP/TAZ signaling pathway, which has been suggested to be involved in primordial follicle activation^27–30^. Based on this, our expectation was that RNA FISH staining of *UHRF1* would be prominent in oocytes residing in morphologically staged primordial follicles, while *CCN2* would be prominent in oocytes within transitioning primordial or primary follicles.

Using the RNA probe for *UHRF1*, we quantified the mRNA puncta within oocytes residing in 8 primordial and 3 transitioning primordial follicles (**Figure 5A,B**). The mRNA puncta counts for *UHRF1* ranged from 53-99 across the 3 transitioning primordial oocytes and 15-147 across the 8 primordial oocytes (**Figure 5B**). With the significant variability in both follicle stages the puncta counts lacked significant difference between them (**Figure 5C**). This is in contrast to single-oocyte RNAseq data, where expression levels of *UHRF1* are significantly different between C1 and C2 (**Figure 5D**).

**Figure 5.**
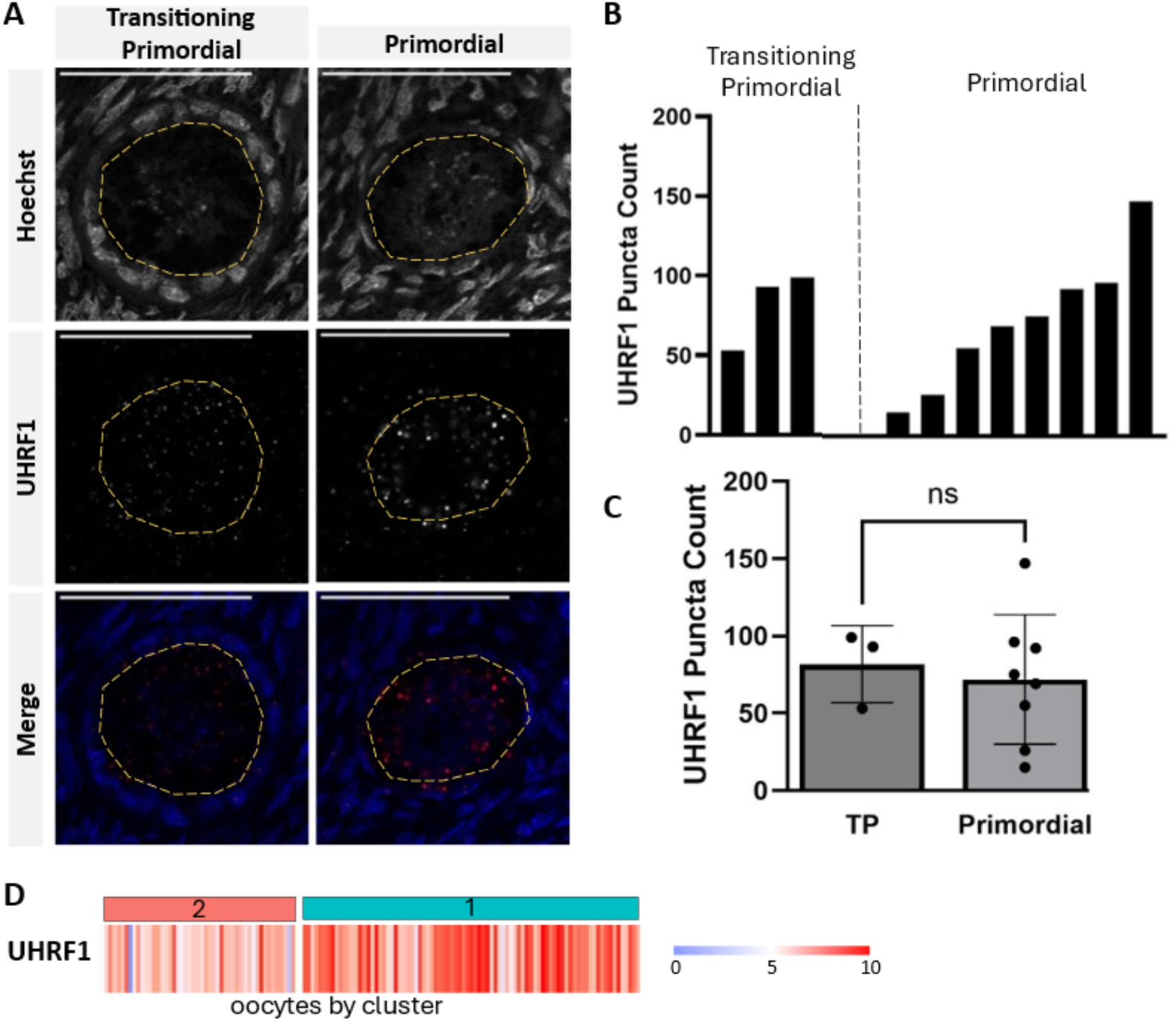
UHRF1 mRNA expression in ovarian tissue to validate the cluster identities. **A.** Representative images of UHRF1 within transitioning primordial and primordial follicles (left-right) and Hoechst, UHRF1, and the merged images (top-middle-bottom). Scale bar: 50 µm. Yellow dashes indicate oocyte boundary. **B.** Bar graph shows UHRF1 mRNA puncta count for all oocytes measured at the transitioning primordial stages and at the primordial stage. **C.** Average UHRF1 puncta counts for transitioning primordial (TP) and primordial stage shows no significant differences. **D.** Heatmap showing UHRF1 expression level for all 133 oocytes by cluster. Color gradient represents logged expression values.

In a similar fashion we quantified the mRNA puncta counts for *CCN2* in the oocytes of primordial, transitioning primordial, and primary follicles and similarly observed a wide range of variability (**Figure 6A,B**). The mRNA puncta counts for *CCN2* ranged from 4 to 43 in 25 transitioning primordial/primary oocytes and from 5 to 47 in 11 primordial oocytes, with no statistical differences between the stages (**Figure 6 B,C**). Once again, the expression levels of *CCN2* in 133 oocytes showed significant C1-C2 difference despite the variability in both clusters (**Figure 6D**).

**Figure 6.**
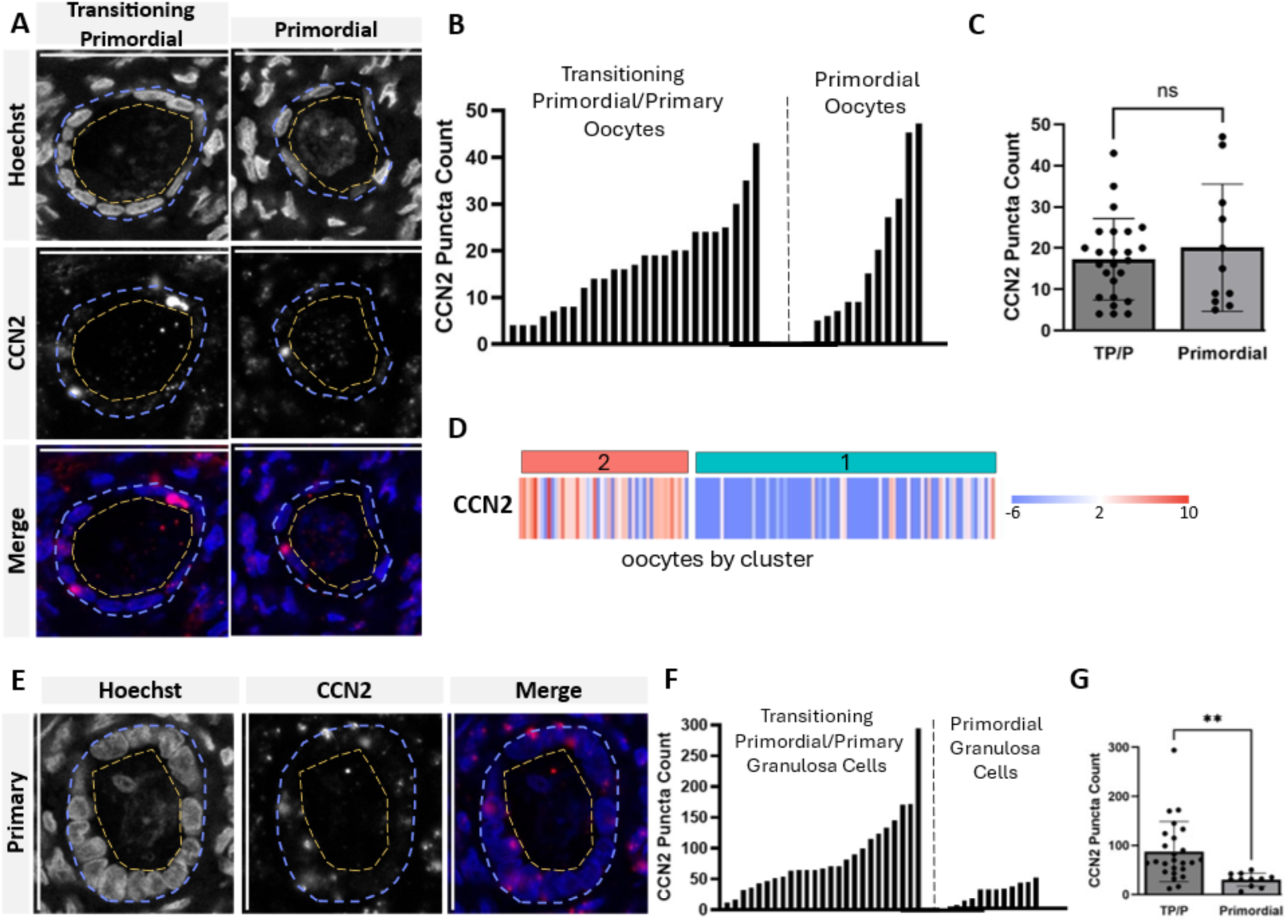
CCN2 mRNA expression in ovarian tissue to validate the cluster identities. **A.** Representative images of CCN2 within transitioning primordial and primordial follicles (left-right) and Hoechst, CCN2, and the merged images (top-middle-bottom). Scale bar: 50 µm. Yellow dashes indicate oocyte boundary. Light blue dashes indicate granulosa cell boundary **B.** Bar graph shows CCN2 mRNA puncta count for all oocytes measured at the transitioning primordial/primary stages combined and at the primordial stage. **C.** Average CCN2 puncta counts for transitioning primordial/primary stages combined (TP/P) and primordial stage shows no significant differences. **D.** Heatmap showing CCN2 expression level for all 133 oocytes by cluster. Color gradient represents logged expression values. **E.** Representative images of CCN2 within primary follicles and Hoechst, CCN2, and merged images (left-middle-right). Scale bar: 50 µm. Yellow dashes indicate oocyte boundary. Light blue dashes indicate granulosa cell boundary. **F.** Bar graph shows CCN2 mRNA puncta count for granulosa cell layer of follicles at the transitioning primoridal/primary stages combined and at the primoridal stage. **G.** Average CCN2 puncta counts for granulosa cells in transitioning primordial/primary stages combined (TP/P) and primordial stage shows significantly higher CCN2 puncta counts in TP/P stages compared with primordial follicles, *p* = 0.005.

Further, *CCN2* data brought an interesting discovery: its mRNA puncta in transitioning primordial/primary follicles are more abundant in the granulosa cell layer than in oocytes (**Figure 6E-G**), with a range of 12-294, and it was significantly higher than the granulosa cells in primordial follicle (range 6-50, *p* = 0.005). In other words, while CCN2 was chosen as a C2 oocyte marker, it is a better marker for granulosa cells in transitioning and primary stages. A recent report by Doherty et al. demonstrated that growing mouse oocytes receive mRNAs from granulosa cells through RNA trafficking along transzonal projections^31^. If similar cytoplasmic shuttling took place in the human samples we analyzed, C2 oocytes could show a higher level of CCN2 even if the difference originated in the surrounding granulosa cells. Additional C1-C2 markers without this complication need to be examined to resolve these possibilities.

## Discussion

Gene expression profiles of human oocytes in the earliest stages of folliculogenesis have been extremely difficult to obtain due to technical limitations in separating oocytes from surrounding somatic cells and the limited availability of human tissue from healthy reproductive-age donors. In this study we overcame these hurdles by analyzing individually denuded oocytes from the ovarian cortex of nine healthy donors. We utilized a recently reported^32^ protocol for dissociation and collection of oocytes and analyzed 133 denuded oocytes. Compared to slide-based laser capture microdissection or analyses of isolated follicles, the unique value of our data lies in the decoupling of the oocyte profile from that of somatic cells.

Since the ovarian cortex tissue we analyzed primarily contained primordial and transitioning primordial follicles (**Figure 1D**), and much fewer primary follicles, we interpret the C1 and C2 oocytes as spanning the primordial-to-transitioning stages, but not further into the primary stage. While the C1-C2 difference aligns with the primordial-primary gene list from Rooda et al. (**Figure 3A-B**), it aligns equally well with the dormant-activated gradient within the primordial follicles (**Figure 4A-B**). The C1-C2 DEGs showed enrichment for similar biological processes (**Figure 1C**) as reported by Rooda et al. In their study, genes higher at the primordial stages are enriched for those linked to cell cycle, chromosomal segregation, and meiosis, while those found higher during follicle growth are linked to ECM and biosynthesis^14^. Our study affirmed that these biological processes are involved in the maturation process of the oocyte, in parallel with the developmental progression of the follicles. Expanding on Rooda et al. primordial-primary and dormant-active genes, our 133-oocyte data yielded new genes of interest. For example, there were several spermatogenesis associated genes (*STPG1, SPATA6,7,20, SPAG4,5,6,9, SLC26A8*) that did not show particularly high fold change in either cluster but showed up abundantly in the oocyte transcriptomes (**Supplemental Data File S1**). These genes could be of interest for future research on oogenesis.

According to the current understanding, the first milestone in oocyte development is the transition from a quiescent state (within primordial follicles) to a fully activated state (within primary follicles). The transitioning primordial follicle state bridges the two, characterized by the presence of a mixed population of squamous and cuboidal granulosa cells. While the C1 and C2 oocytes seem to correspond to those found in primordial and transitioning follicles, our attempts to validate the RNA markers in situ have so far been inconclusive. We have chosen two markers from the C1-C2 DE genes and have not been able to observe a clear difference in their RNA puncta counts between morphologically staged follicles (**Figure 5 and Figure 6**). There are multiple possible explanations. The first is that additional markers need to be tested, and preferably in a larger number of follicles in more samples. This way, subtle differences may become statistically significant.

The second possibility is related to the ongoing uncertainty as to whether oocytes in apparently primordial, transitioning primordial, and primary follicles show a strictly ordered, stepwise transition of transcriptional patterns that parallels the shift of the follicle stages. Follicles in Rooda et al. were staged by granulosa cell morphology^14^, and the authors noted that morphology of granulosa cell alone was not enough to distinguish follicle stages. In our attempt to validate two DE genes, *UHRF1* and *CCN2*, the *in situ* RNA FISH signals did not distinguish between primordial from transitioning follicles. The discrepancy between (1) RNA expression in isolated oocytes and (2) RNA puncta counts in situ for oocytes residing in staged follicles, if replicated in higher powered studies in the future, could suggest that the oocyte contained in a follicle may have progressed further along, or have lagged behind, the follicle stage based on granulosa cell morphology. This would lead to a shift in our practice of staging follicles based solely on morphology. Should follicles of a given stage contain diverse oocytes with overlapping transcriptomic states, an updated metric, such as the transcriptional makeup of oocytes, would serve as a more accurate indicator of their true maturation stage. More accurate ascertainment of the oocyte’s innate stage will facilitate a better understanding of the interplay between the oocyte and granulosa cells during follicle activation.

Furthermore, the design of *in situ* markers in future studies needs to consider the potential cytoplasmic sharing between oocyte and granulosa cells. A recent study provided evidence that mRNAs from granulosa cells are transported into growing mouse oocytes^31^, and their detection in the oocyte may differ between scRNAseq and RNA FISH.

Ultimately, the oocyte-granulosa cell interplay is best studied with spatial analyses. A recent effort from our group used GeoMx^TM^ spatial transcriptomics to analyze whole sections of human ovaries ^34^. However, GeoMx^TM^ did not have the spatial resolution to resolve the transcriptome of individual oocytes. Instead, we stained the tissue sections with the DAZL antibody that selectively binds to the oocytes at primordial, primary and secondary stages, and analyzed mRNA from DAZL+ regions of interest (ROIs) which likely contain mostly primordial and primary oocytes. These oocyte ROIs clustered together, and revealed four new previously unreported marker genes – *PADI6, UCHL1, ZFAND2A*, and *REC114*. These genes were also present in the scRNAseq set from 133 oocytes reported here validating the spatial data (**Supplemental Data File S1**). In general, it is expected that high-content imaging technologies such as CosMx^TM^, Xenium^TM^, and MERFISH would allow us to measure hundreds to thousands of transcripts spatially, for hundreds of follicles that cover a variety of stages. Such data will support much more definitive analyses of how oocytes and their surrounding support cells coordinate their maturation process^35^. The C1-C2 DE genes we report here will inform the best design of spatial analysis gene panels, and we anticipate rapid advances in coming years towards a detailed understanding of folliculogenesis, which will stimulate critical development in *in vitro* activation as potential therapy.

## Acknowledgements

We thank the members of the Shikanov and Li laboratories for scientific discussions and manuscript feedback. We would also like to acknowledge the University of Michigan’s Advanced Genomics Core, the Histology Core at the University of Michigan School of Dentistry, and the ULAM Pathology Core for RNA processing, histological processing, and whole slide scanning respectively. We thank all human organ donors provided by our partnership with the International Institute for the Advancement of Medicine (IIAM).

---

**Supplemental Figure 1.**
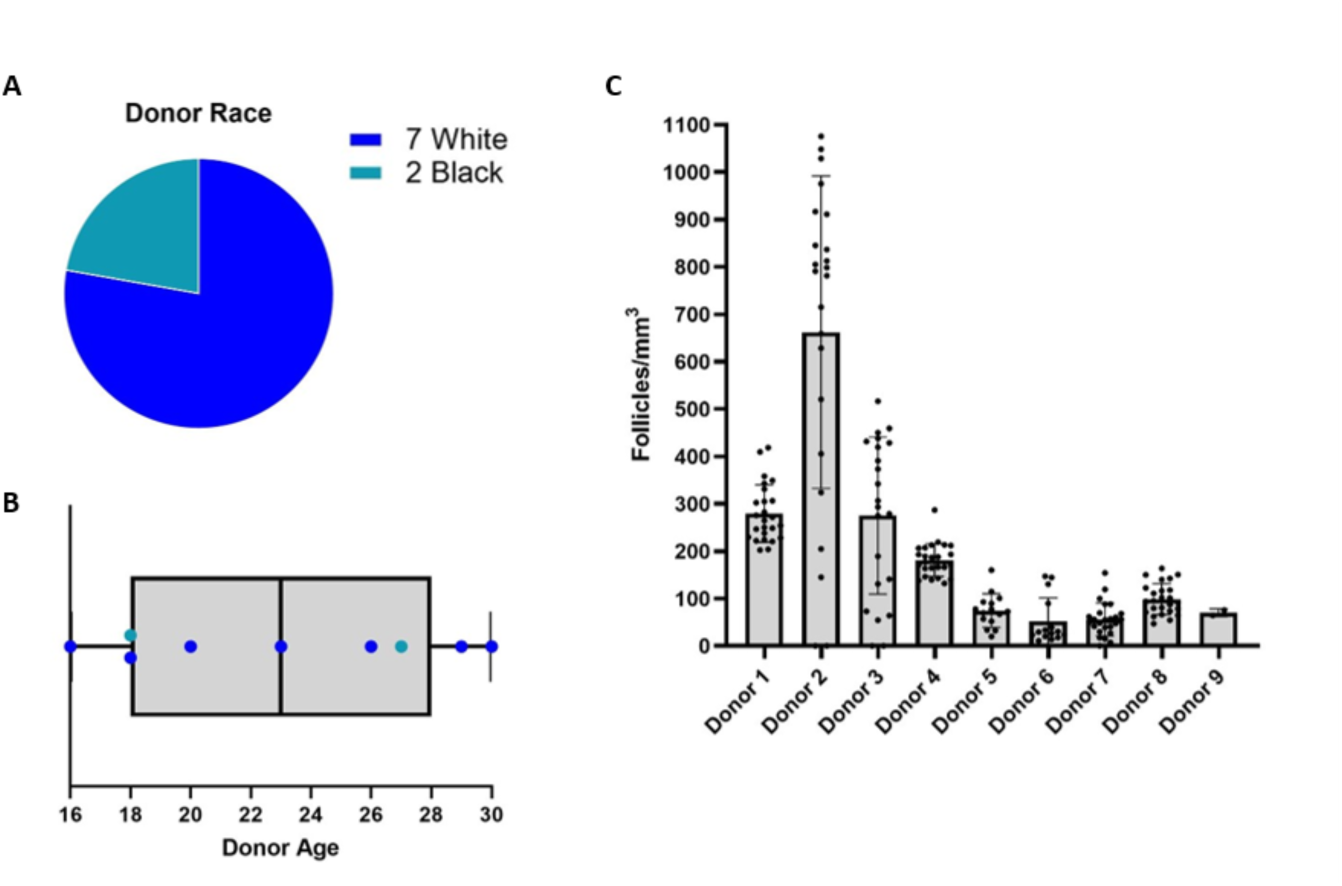
Donor demographics and histological analysis. **A.** Breakdown of reported race for 9 donors used for sequencing. **B.** Spread of donor ages with reported race indicated by teal or dark blue colors. **C.** Follicle densities, as counts per mm^3^, compared across the 9 donors.

**Supplemental Figure 2.**
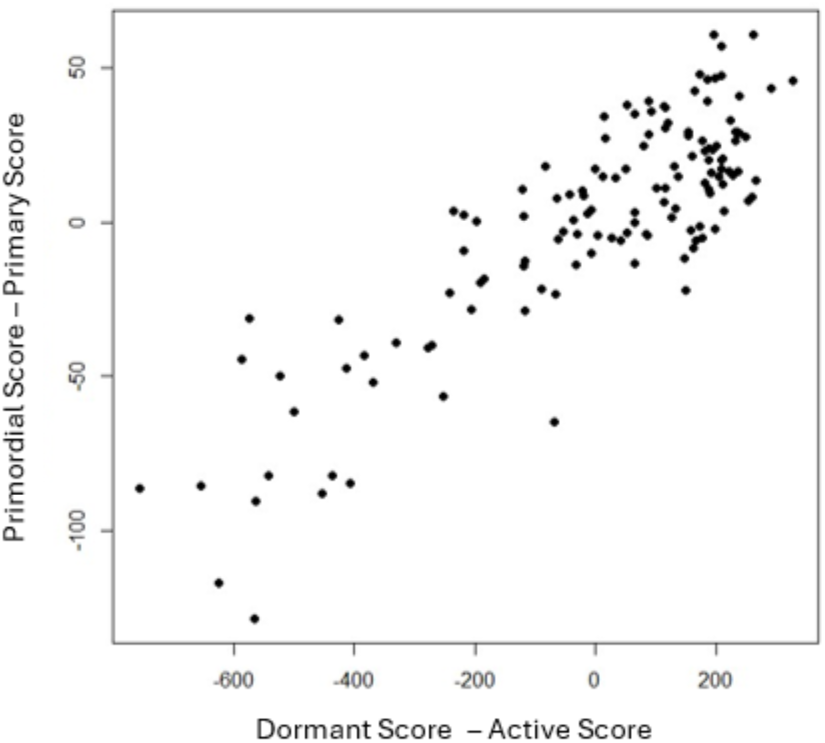
**Concordance between the primordial-primary scores (y axis) and dormant-active scores (x axis) over the 133 oocytes.**

**Supplemental Figure 3.**
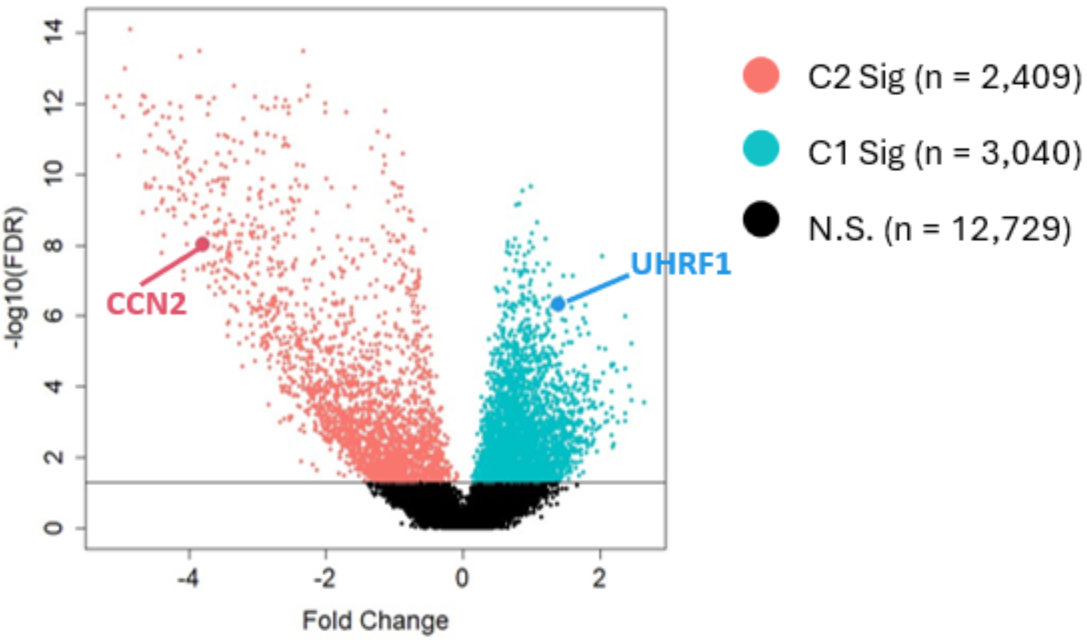
**Selection of UHRF1 and CCN2 as markers for Cluster 1 and 2, respectively, based on DE analysis results.**

**Supplemental Table 1.** Donor metadata, including age, BMI, reported race and Cold Ischemic Time for the sample used in the RNA FISH analyses.

| Supplemental Table 1. Demographics for RNA FISH Donor |  |  |  |  |
| --- | --- | --- | --- | --- |
| Donor # | Age | BMI | Race | Cold Ischemic Time (hr) |
| 10 | 23 | 50.7 | White | 23.6 |

**Supplemental Table 2.** Histological follicle count summary. The number of follicles counted for the 9 donors sequenced at each follicle stage, along with the percent distribution across stages, shown in the parentheses.

| Supplemental Table 2. Histological Follicle Count Summary |  |  |  |  |  |  |  |  |
| --- | --- | --- | --- | --- | --- | --- | --- | --- |
| Donor | Follicle Counts (%) |  |  |  |  |  |  | Total |
| # | Primordial | Transitional Primordial | Primary | Transitional Primary | Secondary | Multilayer | Antral |  |
| 1 | 320 (48.9) | 326 (49.8) | 5 (0.8) | 2 (0.3) | 2 (0.3) | 0 (0) | 0 (0) | 655 |
| 2 | 383 (49.4) | 389 (50.2) | 1 (0.1) | 2 (0.3) | 0 (0) | 0 (0) | 0 (0) | 775 |
| 3 | 362 (36.6) | 529 (53.4) | 52 (5.3) | 27 (2.7) | 8 (0.8) | 10 (1.0) | 2 (0.2) | 990 |
| 4 | 413 (40.3) | 583 (56.8) | 25 (2.4) | 5 (0.5) | 0 (0) | 0 (0) | 0 (0) | 1026 |
| 5 | 35 (30.4) | 78 (67.8) | 1 (0.9) | 1 (0.9) | 0 (0) | 0 (0) | 0 (0) | 115 |
| 6 | 13 (18.8) | 50 (72.5) | 2 (2.9) | 4 (5.8) | 0 (0) | 0 (0) | 0 (0) | 69 |
| 7 | 100 (58.5) | 65 (38.0) | 6 (3.5) | 0 (0) | 0 (0) | 0 (0) | 0 (0) | 171 |
| 8 | 249 (44.0) | 306 (54.1) | 7 (1.2) | 1 (0.2) | 2 (0.4) | 1 (0.2) | 0 (0) | 566 |
| 9 | 92 (43.6) | 100 (47.4) | 13 (6.2) | 4 (1.9) | 2 (0.9) | 0 (0) | 0 (0) | 211 |

